# N-dihydrogalactochitosan reduces mortality in a lethal mouse model of SARS-CoV-2

**DOI:** 10.1101/2021.08.10.455872

**Authors:** Christopher M. Weiss, Hongwei Liu, Erin E. Ball, Ashley R. Hoover, Talia S. Wong, Chun Fung Wong, Samuel Lam, Tomas Hode, M. Kevin Keel, Richard M. Levenson, Wei R. Chen, Lark L. Coffey

## Abstract

The rapid emergence and global dissemination of SARS-CoV-2 that causes COVID-19 continues to cause an unprecedented global health burden resulting in nearly 7 million deaths. While multiple vaccine countermeasures have been approved for emergency use, additional treatments are still needed due to sluggish vaccine rollout, vaccine hesitancy, and inefficient vaccine-mediated protection. Immunoadjuvant compounds delivered intranasally can guide non-specific innate immune responses during the critical early stages of viral replication, reducing morbidity and mortality. N- dihydro**g**alacto**c**hitosan (GC) is a novel mucoadhesive immunostimulatory polymer of β- 0-4-linked N-acetylglucosamine that is solubilized by the conjugation of galactose glycans with current applications as a cancer immunotherapeutic. We tested GC as a potential countermeasure for COVID-19. GC was well-tolerated and did not produce histopathologic lesions in the mouse lung. GC administered intranasally before and after SARS-CoV-2 exposure diminished morbidity and mortality in humanized ACE2 receptor expressing mice by up to 75% and reduced infectious virus levels in the upper airway. Fluorescent labeling of GC shows that it is confined to the lumen or superficial mucosa of the nasal cavity, without involvement of adjacent or deeper tissues. Our findings demonstrate a new application for soluble immunoadjuvants such as GC for preventing disease associated with SARS-CoV-2 and may be particularly attractive to persons who are needle-averse.

**IMPORTANCE:** The ongoing COVID-19 pandemic necessitates new approaches to reduce disease caused by SARS-CoV-2. We tested the immunoadjuvant N-dihydro**g**alacto**c**hitosan (GC), used previously as an immunostimulant for tumor therapy and adjuvant for viral vaccines, as a potential COVID-19 countermeasure. When GC was administered before and after inoculation of a lethal dose of SARS-CoV-2 into the nose of humanized mice expressing an entry receptor for the virus, fewer mice showed weight loss and died compared to mice that received only the vehicle but no GC. GC-treated mice also had lower levels of infectious SARS-CoV-2 in their upper airway. These results suggest that GC may be a candidate to prevent or treat COVID-19.

**Single Sentence Summary:** The immunoadjuvant N-dihydrogalactochitosan diminishes SARS-CoV-2 disease in humanized ACE2 mice, representing a new countermeasure against COVID-19.

## INTRODUCTION

Severe acute respiratory syndrome-like coronavirus 2 (SARS-CoV-2) that was first identified from a cluster of viral pneumonia cases in Wuhan, China in December 2019 [1] has caused more than 670 million cases and nearly 6.9 million deaths globally as of March 2023[2]. Clinical manifestations of 2019 novel coronavirus disease (COVID-19) caused by SARS-CoV-2 typically include fever, non-productive cough, and mild to moderate dyspnea, with severe cases developing pneumonia and acute respiratory distress syndrome [1,3,4]. SARS-CoV-2 morbidity and mortality increase with age and systemic proinflammatory and cardiovascular co-morbidities [1,3,5]. Recovered patients may also exhibit long-duration symptoms including disruption to sensations of taste and smell, and cognitive impairment (colloquially referred to as “brain fog”) resulting from neurological involvement [6, 7].

Like SARS-CoV, SARS-CoV-2 uses the angiotensin converting enzyme 2 (ACE2) for cell entry [8, 9]. Small animals including mice, hamsters, and ferrets, have been fundamental in defining SARS-CoV-2 pathogenesis and developing medical countermeasures [10, 11]. Transgenic mice expressing the human ACE2 (hACE2) gene from the human cytokeratin 18 (K18) promoter have served as an especially useful model of severe disease due to well characterized genetics and ease of use [12–14]. Intranasal inoculation of K18-hACE2 mice (hereafter termed hACE2) with SARS-CoV-2 results in a high viral burden in the lungs, which diminishes past day 7, and elevated viral loads detectible in the brain at day 7 [13].

SARS-CoV-2, which can be transmitted in small-droplet aerosols from person to person, can take up to 14 days to produce symptoms [15]. Public health countermeasures have evolved as additional treatments and information regarding transmission risks for COVID-19 have become available. In addition to face coverings and physical distancing, multiple vaccine candidates and therapeutic interventions have now received emergency use authorization by the United States Food and Drug Administration (FDA) [16]. While ‘herd immunity’ through vaccination remains the target, it is estimated that 70-90% of the global population must be immune in order to interrupt transmission [17]. This target becomes even more challenging with high levels of vaccine skepticism, slow production, inequitable distribution, and the rise of variants of concern capable of more efficient transmission or vaccine escape. Pre- and post-exposure countermeasures can help fill the vaccine gap by reducing COVID-19 morbidity and mortality in unprotected communities and thus reduce the global burden of the pandemic. The nucleoside analog, remdesivir [18], is the only current FDA-approved therapeutic approved for non-hospitalized COVID-19 patients and was authorized based on its ability to shorten recovery time [19], but it is not ideal for clinical use due to its only moderate clinical efficacy [20], as well as its requirement for intravenous administration. Although there are many active trials of therapeutic agents with more than 10 drugs or biological products holding emergency use authorization, including Nirmatrelvir/Ritonavir (Paxlovid) and Molnupiravir (Lagevrio), most are indicated for patients with severe COVID-19 [21, 22]. Despite these advances, there are presently no drugs available for high-risk exposure use to protect against SARS-CoV-2. To circumvent this gap, we repurposed an immunostimulant used for cancer immunotherapy and as a vaccine adjuvant with a goal of mitigating COVID-19 disease.

The parent compound of GC, chitosan, is a linear biological polysaccharide polymer of β-0-4-linked N-acetylglucosamine and is produced from alkaline treatment of the chitin exoskeleton of crustaceans. Chitosan is approved by the FDA for tissue engineering and drug delivery. Chitosan shows broad-acting antiviral [23] and immunoadjuvant properties [24, 25] including interferon (IFN) induction [26, 27] that is critical for viral control [28–30], and has been evaluated as an antiviral therapy for respiratory viruses [31–33]. Chitosan and various derivatives have been explored as COVID-19 treatments and vaccine adjuvants [34–39]. Modification of chitosan by covalently attaching galactose sugars to the free amino acids on the polysaccharide backbone produces N-dihydro**g**alacto**c**hitosan (GC) [40, 41]. GC was initially designed to further improve immune-stimulating functionality of the molecular backbone by adding glycan moieties that can bind to C-type lectin receptors on antigen presenting cells [42] and lead to a downstream immune response while retaining the mucoadhesive properties of the parent molecule [43]. Furthermore, GC has improved solubility at physiological pH ranges compared to its parent molecule, and hence is more suitable as an injectable agent. With these modifications, GC has been developed for use in human interventional immuno-oncology to stimulate systemic anti-tumor immunity [44]. GC recruits granulocytes at the injection site [45] and stimulates activation of dendritic cells and macrophages through upregulation of co-stimulatory molecules CD40, CD86, and MHCII *in vitro* and *in vivo* [46, 47] . Macrophages also show increased nitric oxide production and phagocytic abilities in response to GC [46]. Single-cell RNA sequencing indicates enrichment of type I IFN signaling in multiple innate immune cells, including monocytes, M1 macrophages, and neutrophils in tumors following GC administration [48]. Our prior work showed that GC mixed with recombinant SARS-COV-2 spike and nucleocapsid antigen vaccines delivered intranasally protects mice from lethal disease after intranasal SARS-CoV-2 inoculation. In that study, when 0.75% GC was delivered intranasally to control mice, CD45^+^ and B cell cellularity increased in cervical lymph nodes accompanied by an influx of migrating and resident dendritic cells and CD4^+^ and CD8^+^ T cells. These data show that GC induces a local immune response that induces lymphocyte migration and activation in the nasal-associated lymphoid tissue (NALT)[49]. The upregulation of innate immune responses and recruitment of cellular responses to the NALT, which is the SARS-CoV-2 exposure site, suggests that broad-acting non-toxic immunoadjuvants such as GC that can be intranasally may modulate SARS-CoV-2 infection. Intranasal instillation may generate a robust immune protection in the upper respiratory tract and may be more widely accepted than vaccine injections. Furthermore, non-specific immunoadjuvants such as GC could offer a benefit over more\ narrowly targeted antivirals by providing protection against a potentially broad range of pathogens, potentially including future agents not yet affecting public health.

In the present study, we explored the use of GC for prophylaxis against SARS-CoV-2 infection applied pre- and post-exposure to lethal dosages of the virus in transgenic hACE2 mice. To simulate the use of solubilized GC as a nasal spray, we applied the compound intranasally twice before exposure and once following exposure of mice to a human isolate of SARS-CoV-2. Our data demonstrate a strong protective effect of GC in preventing mortality in this lethal model of disease. Fluorescent-labeled GC administered intranasally remained in the nasal cavity. Pre- and post-exposure countermeasures may help fill the vaccine gap by reducing COVID-19 morbidity and mortality and reduce the global burden of the pandemic.

## RESULTS

### N-dihydrogalactochitosan reduces SARS-CoV-2-associated mortality in hACE2 transgenic mice

To determine the antiviral efficacy of N-dihydro**g**alacto**c**hitosan (GC) against SARS-CoV-2, 6-week-old hACE2 [B6.Cg-Tg(K18-ACE2)2Prlmn/J] mice were treated intranasally (i.n.) with 0.75% GC in neutral buffered saline 3 and 1 days prior to virus exposure and then 1 day post exposure (**Fig. 1a**). To ensure that the dose, schedule, and route of GC administration were well tolerated, pilot studies using 14 additional male and female hACE2 mice showed that GC treatment alone does not produce adverse clinical signs or pulmonary lesions. Lung histologic scores in mice administered GC were not significantly different from mice treated with PBS (**Supplementary Fig. 1, Supplementary Table 1**). Mice were inoculated i.n. 1 day prior to the final GC treatment with 10^3^ or 10^4^ plaque forming units (PFU) of SARS-CoV-2, a route and dose intended to produce high lethality in this model. Inocula were back-titrated to verify the administered dose. Tracheal swabs were collected for the first 3 days post inoculation and animals were monitored daily until day 14 for weight loss and health status.

**Fig. 1:**
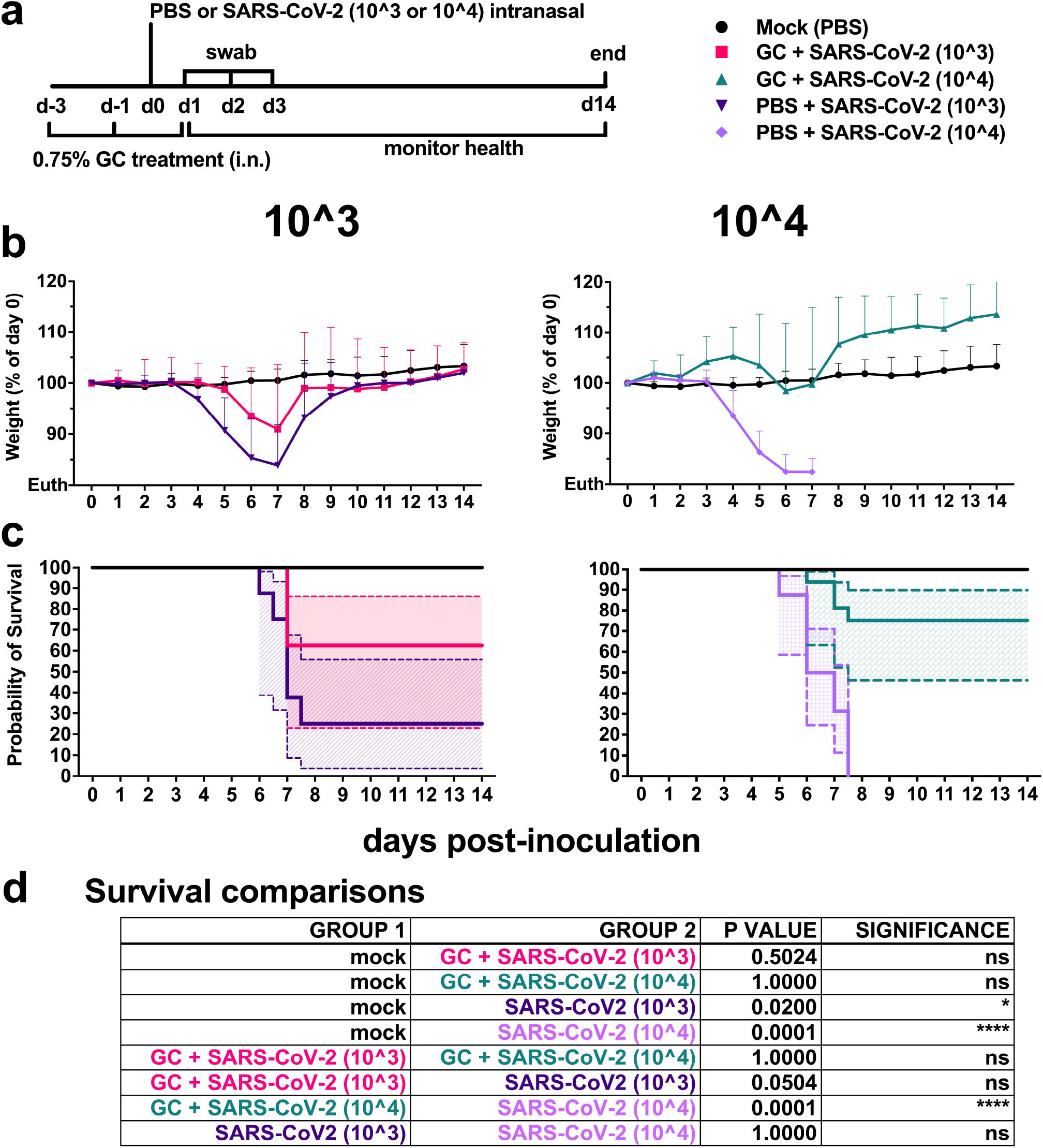
N-dihydrogalactochitosan protects mice from SARS-CoV-2 mortality. **(a)** Experimental design where 6-week-old male and female hACE2 transgenic mice were treated with 0.75% GC or PBS delivery vehicle at days -3, -1 and +1 post-inoculation. Mice were inoculated at day 0 with 103 or 104 PFU of SARS-CoV-2 or PBS (mock). Animals were weighed daily, and throats were swabbed on days 1 through 3. (**b**) **Weight** represented as a percentage of individual mouse weight at the time of inoculation. A main effect only model two-way ANOVA with Tukey corrected multiple comparisons yielded an *F*=52.80, *p*<0.0001, 4 degrees of freedom. Each symbol represents an individual mouse, and the horizontal lines show geometric mean and error bars are geometric standard deviation. **(c) Survival proportions.** The solid lines are survival proportions, and the shaded boxes show 95% confidence intervals. **(d) Logrank Mantel-Cox comparisons of survival proportions** with Bonferroni corrected p-values, multiple pairwise tests, 1 degree of freedom. n=8-16 per group, 2 combined experiments.

Weight remained stable for the first 3 days in all groups. Weight in infected mice that were not GC-treated rapidly declined, starting at day 4 in the 10^4^ PFU group, and at day 5 in the 10^3^ PFU groups (**Fig. 1b**). Half of the GC-treated mice administered 10^4^ PFU experienced no weight loss throughout the duration of the study (**Supplementary Fig. 2**); conversely, their growth outpaced mock-infected counterparts treated with saline alone (*p*<0.0001). This effect was not limited to one biological sex and did not correlate with mouse starting weight at the time of study initiation. Additional mice lost weight starting at day 4 but did not reach euthanasia criteria (loss of 20% of initial body weight) and recovered to starting weight by day 10 post-inoculation. Relative to the 10^4^ group, mice inoculated with 10^3^ PFU SARS-CoV-2 exhibited delayed or transient weight loss, with 2 mice experiencing no weight loss over the duration of the study. At the higher inoculation dose of 10^4^ PFU, GC significantly reduced weight loss versus delivery vehicle treated controls (*p*<0.0001). At the lower 10^3^ PFU inoculation dose, GC trended toward protection from weight loss, which was confounded by non-uniform disease in delivery vehicle treated controls (*p*=0.11). Mice treated with delivery vehicle and inoculated with 10^3^ or 10^4^ PFU had a median survival time of 7±3.3 and 6.5±0.9 days, respectively (**Fig. 1c**). At the lower inoculation dose of 10^3^ PFU, GC had an efficacy of 37.5% protection from mortality (*p*=0.05) (**Fig. 1c**). At the higher inoculation dose of 10^4^ PFU, GC reduced SARS-CoV-2 mortality by 75% (*p*<0.0001) (**Fig. 1d**). Together, these results demonstrate the potent efficacy of GC in preventing fatal SARS-CoV-2 disease in transgenic mice.

### N-dihydrogalactochitosan reduces SARS-CoV-2 viral levels in the respiratory tract

We next sought to determine whether GC reduced viral levels in addition to protecting mice from mortality. Infectious SARS-CoV-2 levels were assessed longitudinally in mice by swabbing throats from day 1 to 3 post-inoculation (**Fig. 2a**). SARS-CoV-2 was detectible in tracheal swabs of infected animals at day 1 post inoculation and most animals had no detectible virus by day 3. Virus levels were elevated in mice inoculated with 10^4^ PFU versus 10^3^ PFU at 1 day post inoculation (*p*=0.05), but differences between inoculation doses were not detected after day 2 (*p*>0.99). GC significantly reduced virus levels in tracheal swabs at day 1 and day 2 post inoculation with 10^4^ PFU (*p*=0.0005 and *p*=0.02, respectively). A similar effect was not observed in mice that received the lower inoculum of 10^3^ PFU. All delivery vehicle-treated mice had detectible virus in tracheal swabs following infection, while 29.2% (7/24) of GC-treated mice had no infectious virus isolated between days 1 and 3. Positive virus detection in tracheal swabs was not correlated with lethal disease in mice (*p*=0.62, Fisher’s exact test). Cumulative virus levels in tracheal swabs were calculated as area under the curve for individual animals (**Fig. 2b**). GC significantly reduced the total shed virus levels in tracheal swabs in mice inoculated with 10^4^ PFU from a geometric mean of 225 to 5.6 PFU (*p*<0.0001) and trended toward a reduction in animals inoculated with 10^3^ PFU from 138 to 15 PFU (*p*=0.08).

**Fig. 2:**
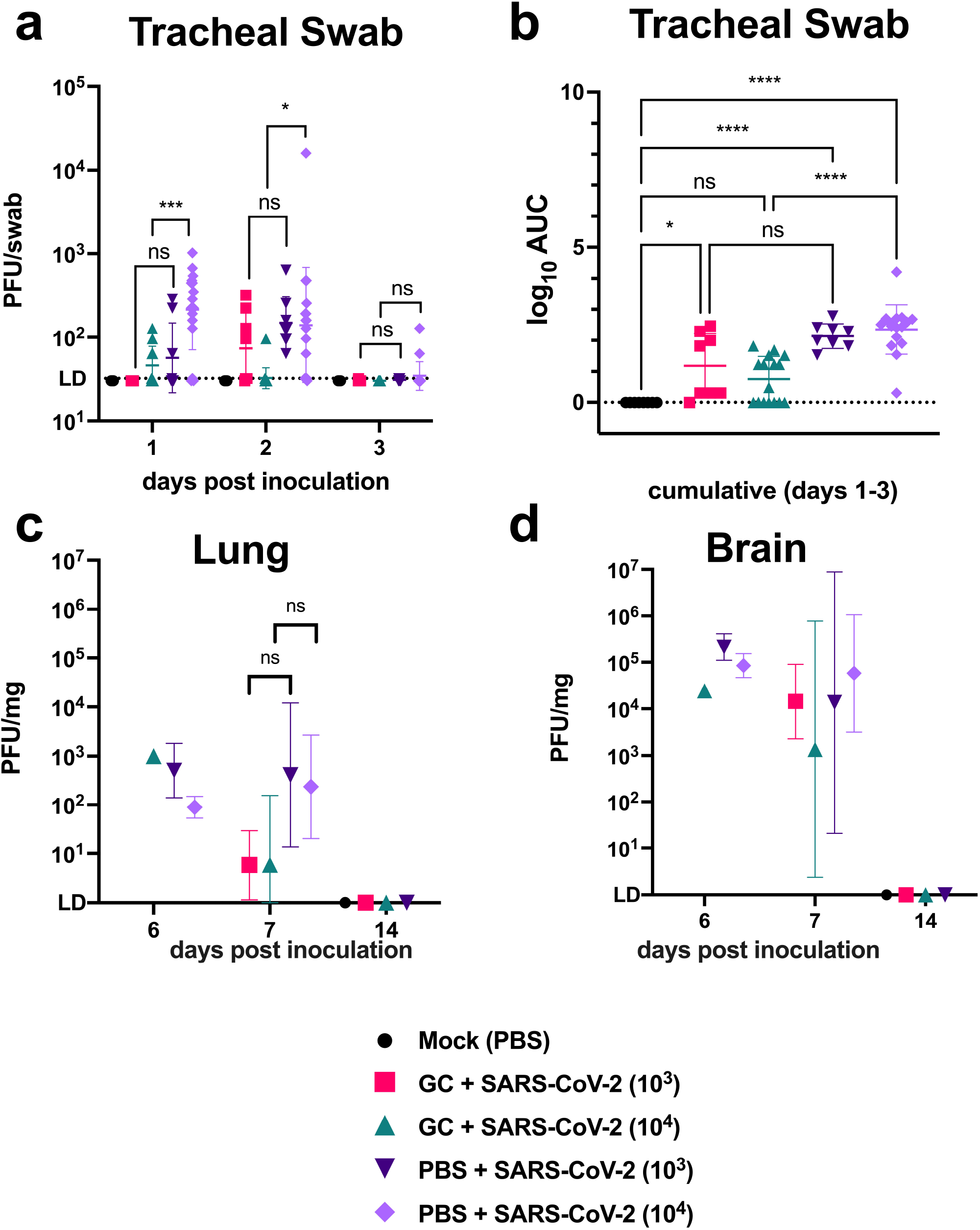
N-dihydrogalactochitosan reduces SARS-CoV-2 detection in the respiratory tract. Infectious SARS-CoV-2 was measured in (**a**) **tracheal swabs** collected from hACE2 transgenic mice on days 1, 2 and 3 post inoculation. Repeated measures two-way ANOVA with Tukey corrected multiple comparisons on log10-transformed values, *F*=16.96, *p*<0.0001, 4 degrees of freedom. (**b) Total area under the curve** shows cumulative virus levels in swabs. ANOVA of log10-transformed total peak area with Tukey corrected multiple comparisons, *F*=7.538, *p*<0.0001, 4 degrees of freedom. Mice were necropsied on days 6, 7 and 14 post-inoculation as euthanasia criteria were met. Infectious virus levels were measured in (**c**) **lung** and (**d**) **brain** at experimental endpoints. Main effects only model two-way ANOVA on log10-transformed values, *F*=2.431, *p*=0.0606, 4 degrees of freedom for lung, *F*=0.8134, *p*=0.5229, 4 degrees of freedom for brain. Symbols are individual animals, horizontal lines show geometric mean, and error bars represent geometric standard deviation. n=8-16 per group, 2 combined experiments. LD = limit of detection, AUC = area under the curve. * *p* < 0.05, *** *p* < 0.001, **** *p* < 0.0001.

As individual animals met humane experimental endpoints, mice were euthanized. Infectious virus levels were measured in the lungs (**Fig. 2c**) and brain (**Fig. 2d**) from mice on days 6, 7, and 14. Although mean virus levels were not significantly different, GC treatment trended toward reducing lung virus levels (*F*=2.431, *p*=0.06, two-way ANOVA) with cumulative effects in animals inoculated with 10^3^ PFU driving the main treatment effect (*p*=0.06).

Comparatively high viral titers were observed in the brain at the time of death, consistent with previous descriptions in this model [13]. Virus was detectible in the brain in all but 2 infected animals at the time of death, indicating neuroinvasion as a likely cause of morbidity. No significant differences in brain virus levels were observed between treatments (*F*=0.8134, *p*=0.52, two-way ANOVA). No infectious virus was detectible in the lungs or brains of animals euthanized at day 14 post inoculation.

### N-dihydrogalactochitosan reduces the severity of histopathologic lesions associated with SARS-CoV-2 infection in lungs

Intranasal inoculation of hACE2 transgenic mice with PBS followed by mock inoculation resulted in normal lung architecture in most mice, although several animals exhibited mild alveolar septal inflammation, likely associated with i.n. administration **(Fig. 3a)**. By contrast, mice inoculated with 10^3^ or 10^4^ PFU SARS-CoV-2 (**Fig. 3c**, representative 10^3^ image shown) exhibited extensive lymphohistiocytic interstitial pneumonia with a peribronchiolar and perivascular distribution and scattered multinucleated syncytial cells 7 days post inoculation. Bronchiolitis, bronchiolar epithelial and alveolar septal necrosis, hemorrhage, fibrin, edema, and vascular endothelial inflammation were also occasionally noted. SARS-CoV-2-infected mice treated with GC showed lesser histopathologic lesions in the lung (**Fig. 3b**, representative 10^3^ image shown) compared to animals that did not receive GC, and inflammation was focally distributed instead of widespread. To quantify histopathologic changes, lung lesion severity was scored using specific criteria (**Supplementary Table 2**). The percent of the lung that was affected in mice euthanized at 6, 7, or 14 days post inoculation as assessed by image analysis software did not differ significantly between SARS-CoV-2 infected mice treated with GC versus those who were not GC treated(**Fig. 3d**). Despite this, GC treatment significantly reduced the mean lung histopathology severity score across all euthanasia days (although not when the results at days 6 and 7 or 14 were analyzed separately [**Fig. 3e**]) for mice dosed with 10^4^ PFU SARS-CoV-2; scores were reduced from 4.6 to 2.5 (ANOVA, *p* < 0.0001) (**Fig. 3f**). Mean scores for mice treated with GC and 10^3^ PFU of SARS-CoV-2 trended towards being lower than for mice who did not receive GC but were not significantly different (p >0.05). Together, these data show that GC reduces SARS-CoV-2-induced disease in the lung of hACE2 mice, although only significantly in mice administered the higher 10^4^ PFU dose of SARS-CoV-2.

**Fig. 3:**
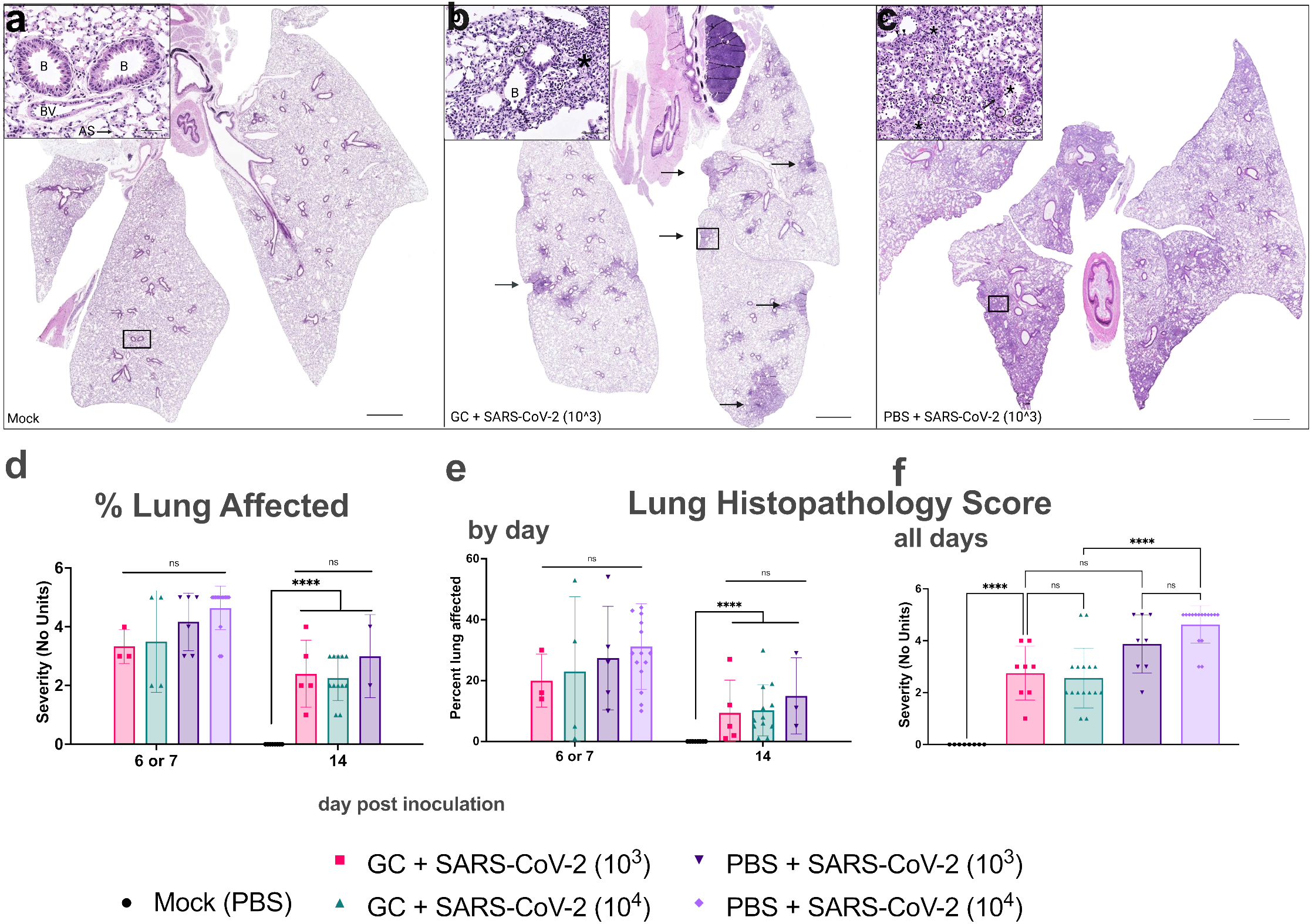
N-dihydrogalactochitosan reduces histopathology associated with SARS-CoV-2 infection in lungs. Lungs from hACE2 transgenic mice were collected at the time of death, thin sectioned, and hematoxylin and eosin stained for histopathological scoring. Images show representative lungs. **(a)** Normal lung from a **PBS-treated, mock inoculated mouse at day 14**. Bronchioles lined by epithelial cells, alveolar septa containing pulmonary capillaries lined by pneumocytes, and a small arteriole are visible (inset). **(b)** Lung from a **GC + 10^3^ PFU SARS-CoV-2 mouse at day 7** with patchy inflammation (black arrows) distributed around airways and affecting approximately 15% of the section. Peribronchiolar alveolar septal inflammation composed primarily of lymphocytes and macrophages (asterisk, inset) with scattered multinucleated cells. **(c)** Lung from a **PBS + 10^3^ PFU SARS-CoV-2 inoculated mouse at day 7** showing widespread, multifocal to coalescing inflammation affecting all lung lobes and approximately 60% of the section. The inset shows lymphohistiocytic bronchoalveolar inflammation (asterisk) with multinucleated syncytial cells (circled), endotheliitis (black arrowheads) and bronchiolar epithelial hyperplasia characterized by disorganization and piling-up of bronchiolar epithelium with increased mitotic figures (black arrows). (**d**) **Total affected lung area** was estimated using image analysis software. **(e-f) Lung lesion severity** was scored according to criteria defined in Supplementary Table 1. Scale bars are 2 mm (subgross) and 50 µm (insets). As=alveolar septa, B=bronchioles, Br=bronchus, BV=blood vessel. ** *p* < 0.01, *** *p* < 0.001, **** *p* < 0.0001, ns=not significant. ANOVA, *F*=35.86, *p*<0.0001, 4 degrees of freedom (e). ANOVA, *F*=10.32, *p*<0.0001, 4 degrees of freedom (e). Symbols are individual animals, bars show the mean, and error bars show the standard deviation, n=8-16 per group, 2 combined experiments.

### Mice treated with N-dihydrogalactochitosan produce neutralizing antibodies after SARS-CoV-2 exposure

We next sought to determine whether mice surviving SARS-CoV-2 inoculation produced a humoral immune response that might protect them from future infection. Serum from blood collected at day 14 post inoculation was assessed for neutralizing antibody against the inoculation strain of SARS-CoV-2 by plaque reduction neutralization test (PRNT) at the 80% neutralization threshold (**Fig. 4**). All mice surviving virus inoculation generated neutralizing antibodies. Mice in the GC group inoculated with 10^3^ or 10^4^ PFU generated neutralizing titers of 1:676 and 1:861 (geometric mean), respectively. In comparison, PBS-administered mice that received an inoculation dose of 10^3^ PFU had a mean neutralizing titer of 1:1448. Neutralizing antibody titers were not different across treatments (p>0.99) or virus dose, although PBS-administered mice inoculated with 10^4^ PFU SARS-CoV-2 were unavailable for comparison due to uniform lethality. These data confirm that surviving mice were infected and indicate that animals were able to develop adaptive immune responses that could potentially protect against reinfection.

**Fig. 4:**
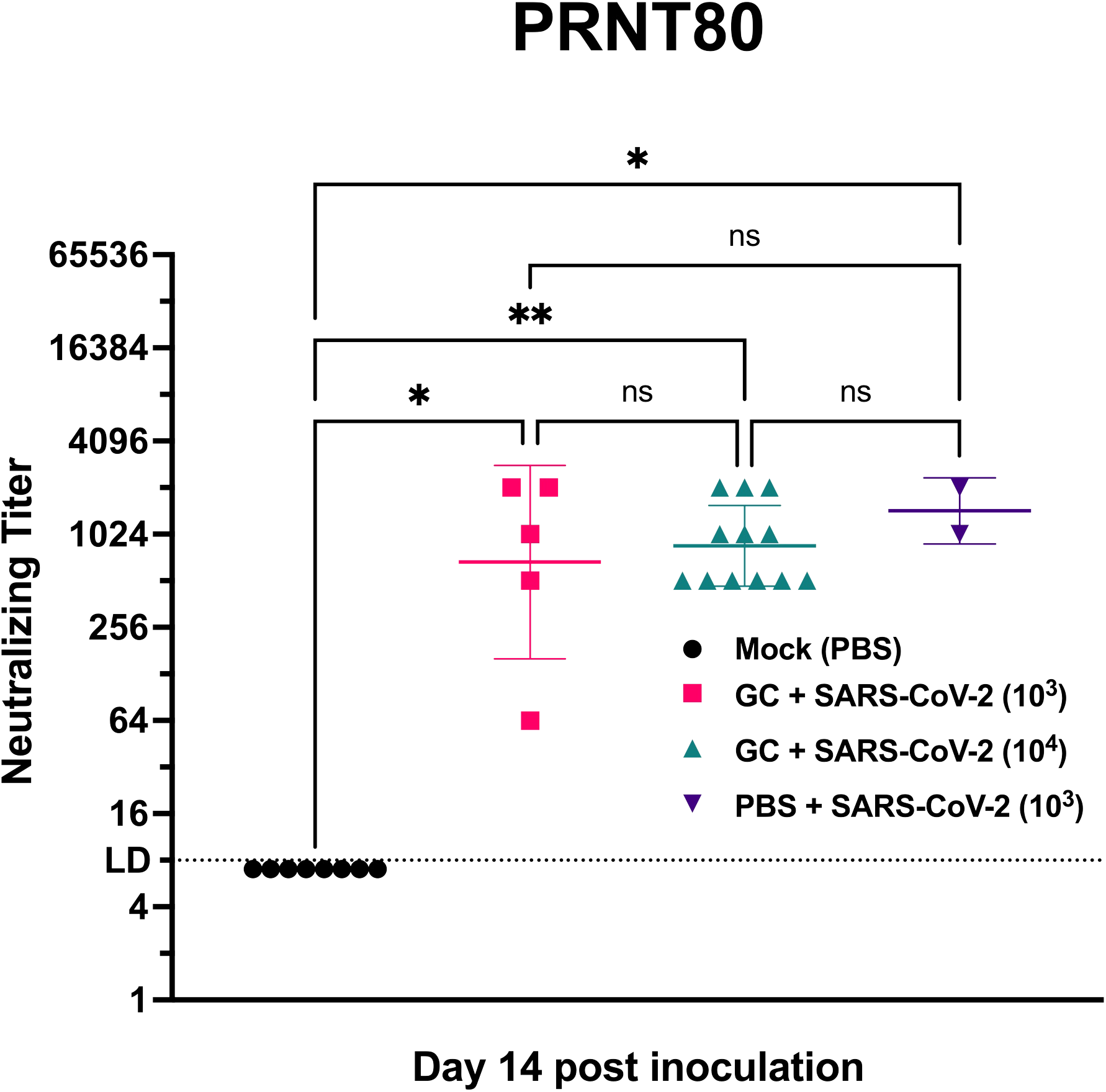
N-dihydrogalactochitosan treated mice surviving SARS-CoV-2 inoculation produce neutralizing antibodies. Neutralizing antibody was assessed by 80% plaque reduction neutralization test (PRNT80) in serum from hACE2 transgenic mice surviving to day 14 post inoculation. No PBS treated mice inoculated with 104 PFU SARS-CoV-2 were available for comparison due to uniform mortality. * *p* < 0.05, ** *p* < 0.01, LD = limit of detection. Kruskal-Wallis test with Dunn corrected multiple comparisons, *H*=17.87, *p*=0.0005, 3 degrees of freedom. Symbols are individual animals, horizontal lines are geometric mean and error bars show geometric standard deviation, n=2-12 per group, two combined experiments.

### Fluorescent labeled N-dihydrogalactochitosan administered intranasally in mice localizes to the mucous of the nasal cavity

Finally, we administered fluorescent labeled GC (FITC-GC) i.n. to mice to examine retention kinetics in the nasal concha. Two mice per time point were FITC-GC treated once and then euthanized 2h, 9h, or 2d later, or were treated twice (2X) and harvested 3d later; for these mice, the second treatment was administered 1 day before euthanasia. Negative control mice were not administered FITC-GC. Histopathology was evaluated and quantified using a scoring rubric developed for the nasal cavity (**Supplemental Table 3**). Fluorescence in the nasal cavity was visualized with microscopy measured by calculating the percent area of the total nasal conchae per slide with a fluorescent signal. Fluorescing aggregates identified within the oral cavity of several treated mice were not included in the measurements, as they were outside the nasal conchae. Negative control mice had no fluorescent foci (**Fig. 5a**). For FITC-GC treated mice, fluorescing foci were predominantly within the lumen or loosely adhered to the nasal mucosa, with infrequent fluorescing areas within the epithelium (**Fig. 5b-f**).

**Fig. 5.**
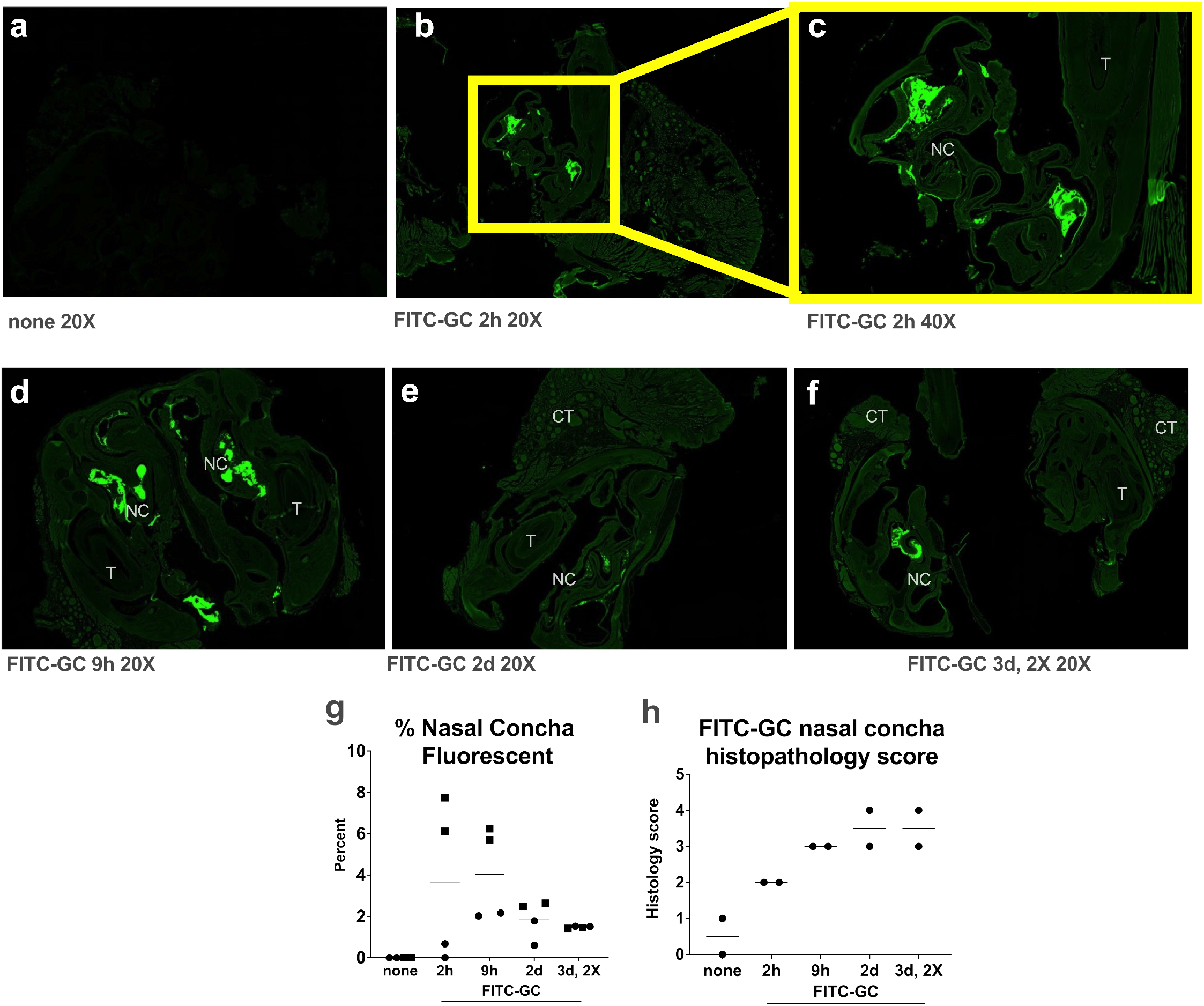
Fluorescent labeled GC (FITC-GC) administered intranasally to mice localizes to the nasal cavity and immune cell influx results in nasal concha histopathology scores. Fluorescent images of nasal concha from mice administered (a) no FITC-GC or (b-f) 0.75% FITC-GC i.n. administered once, and euthanized 2h, 9h, 2d or 3d later. Mice in the group euthanized 3d after the first treatment were administered a second treatment 1d before euthanasia. Panel (c) is at a magnification of 20, (b-f) are at a magnification of 20X. NC is nasal conchae, T is tooth, CT is connective tissue. (g) Fluorescence measured as percent of total image area, limited to nasal mucosa. Matching symbols for each time show 2 measurements from the same mouse. (h) Nasal concha histopathology scores based on the scoring rubric in Supplemental Table 2. N=2 mice per time. The small group size precluded statistical assessments.

Although small group sizes preclude statistical assessments, the fluorescent signal of FITC-GC in the nasal cavity of all but 1 FITC-GC treated mouse appeared higher than in negative control mice (**Fig. 5g**). Histologically, fluorescent material was typically associated with inflammatory cells, mucous, and debris. Histology scoring in the nasal concha of these mice showed that while PBS treated mice had low scores, FITC-GC treatment increased scores (**Fig. 5h**). Treatment of the group of mice euthanized 3d post treatment with 2 administrations of FITC-GC did not increase the percent nasal concha fluorescent or histology scores. All FITC-GC treated mice showed moderate rhinitis characterized by mixed inflammatory mucosal or submucosal infiltration and or nasal cavity exudate composed of neutrophils, macrophages and lymphocytes and mild to moderate hemorrhage, fibrin, edema, and sometimes necrotic debris. Together, these data support a process whereby GC administered i.n. in mice remains confined to the superficial mucosa and acts as a mild irritant without eliciting deeper inflammation.

## DISCUSSION

Intranasal administration of GC to prevent or treat SARS-CoV-2 represents a novel application of this potent immunostimulatory compound. In the present study, GC was well-tolerated and prevented lethal disease in up to 75% of treated humanized mice, while all mice not receiving GC but inoculated with 10^4^ PFU SARS-CoV-2 required euthanasia. All surviving animals had neutralizing antibodies detectable at 14 days post inoculation. Similar SARS-CoV-2 antibody titers were detected in GC-treated and untreated animals. Mice that received GC treatment also displayed lower levels of infectious SARS-CoV-2 in tracheal swabs collected 1 to 2 days post inoculation. Reducing virus levels in the upper airway has gained particular significance as recent vaccine breakthrough cases with variants of concern demonstrate SARS-CoV-2 shedding at similar levels to unvaccinated individuals, which is a key determinant of transmission potential [50]. Our findings demonstrate a reduction in viral burden and virus-induced disease along with a resultant increase in survival due to GC treatment.

GC is a polymeric mixture containing strands of varying lengths of selectively galactose-conjugated and partially deacetylated β-0-4-linked N-acetylglucosamine molecules with a specific range of molecular weights [40,41,44,45]. Synthesized and purified under Good Manufacturing Practice (GMP) conditions, N- dihydrogalactochitosan is a well-characterized variant of GC. Characterization and quality control testing ruled out contamination from endotoxins, heavy metals, and other impurities, which eliminates a major confounding factor in the research of these naturally derived molecules, and of chitosan in particular [51]. Water solubility, immunological properties, biocompatibility, and a favorable toxicity profile are key features of GC. The unconjugated base polymer, chitin, is a primary structural component of cell walls for organisms ranging from fungi to arthropods [23]. Chitosan, a derivative of chitin, is deacetylated through alkaline treatment and is marketed as a nutritional supplement [52] and used as a biopolymer [53]. Unmodified chitosan has been used successfully to treat influenza A virus infection in mice [31], demonstrating a potential application as a non-specific antiviral compound for respiratory virus infection. However, unmodified chitosan has low solubility in neutral buffered aqueous solution and requires acidic formulation; poor characterization, purification and lack of controlled synthetic pathways lead to poor reproducibility and unpredictable outcomes [51]. GC circumvents these limitations through a controlled and reproducible process of synthetically attaching galactose to the free-amino groups of the chitosan base, improving solubility (and thus bioavailability) while maintaining a physiological pH. GC has been previously used as a combination anti-tumor therapy due to its immunoadjuvant properties [46, 48], but its utility as a broad-acting antiviral compound has not previously been investigated.

Immunoadjuvants stimulate non-specific innate immune responses through various mechanisms. Some use pattern-recognition receptors, including the Toll-like receptor (TLR) family of antigen detection complexes. While the mechanism of SARS-CoV-2 protection has not been determined for GC, chitosan, and similar polymers of N-acetylglucosamine, interact with TLR2, which serves as a rationale for inclusion of chitosan as a vaccine adjuvant. TLR2 signals through myeloid differentiation factor 88 (MyD88) to stimulate the nuclear factor kappa B (NF-κB) pathway and downstream inflammatory and anti-microbial cytokine responses [54, 55]. TLR engagement of the canonical NF-κB pathway up-regulates both tumor necrosis factor alpha (TNF-α) and interleukin-6 (IL-6), two potent pro-inflammatory effectors. IL-6, which is necessary for antiviral immunity against other virus families, has been identified as a target of dysregulation associated with hyperinflammatory responses in the lungs during SARS-family coronavirus infections. SARS-CoV-1 nucleocapsid protein can stimulate NF-κB activation and IL-6 production independent of infection [56]. Prolonged high levels of IL- 6 are correlated with severe COVID-19 outcomes in humans, and a similar association has been found in ferrets infected with SARS-CoV-2 [57]. The timing of IL-6 production may play a critical role in determining whether viral clearance is achieved or alternatively, hyperinflammatory responses result. Early, but not late, induction of IL-6 during respiratory syncytial virus (RSV), influenza A, and rhinovirus infection promote viral clearance and limit prolonged inflammation by establishing regulatory T cell populations that limit virus spread [58]. Similarly, timed induction of pro-inflammatory responses by GC treatment pre- and post-infection may promote viral clearance before virus-mediated dysregulation of this pathway can occur. As such, future studies should explore the timing of GC treatment and regulation of IL-6 and associated pro-inflammatory pathways.

The data in this study demonstrate that treatment with GC does not impact development of a neutralizing antibody response by day 14 after SARS-CoV-2 inoculation, even though it reduced infectious SARS-CoV-2 in tracheal swabs. These data suggest GC functions as an immunoadjuvant that can modulate innate immune responses after SARS-CoV-2 infection, leading to stimulation of robust antiviral adaptive immunity. Experimental evidence from cell culture and animal models supports at least four mechanisms by which complex carbohydrates such as GC could stimulate the innate and adaptive immune systems [59–62]. First, GC recognition by sensors on macrophages and dendritic cells (DC) can stimulate innate immune defenses [63–65].

While immune receptors for GC have not been identified, chitosan binding to C-type lectin receptors such as dectin-1 can initiate innate immune signaling [66, 67]. Second, GC-mediated antigen uptake [68] and antigen presentation by DCs could lead to CD4 and CD8 T cell responses [45, 46]. Studies in mice show GC induces type 1 IFN [48], leading to enhanced DC activation and robust CD4 T helper cells [60]. Activation of the STING pathway by type I IFN is a known important antiviral pathway independent of neutralizing antibodies[49]. Notably, DNA sensor activation is mediated by chitosan-induced cellular DNA release. Third, physical antigen sequestration and slow antigen release within draining lymph nodes facilitated by GC could potentially prolong germinal center reactions, thereby enhancing humoral immunity by fostering affinity maturation of B cells [69, 70]. Fourth, GC triggers recruitment of neutrophils, macrophages, and lymphocytes in the nasal cavity indicating a heightened innate immune response that could eliminate SARS-CoV-2 (and other respiratory pathogens) and infected cells at the entry site via mechanisms such as phagocytosis, formation of neutrophil extracellular traps, the release of reactive oxygen species, and inflammatory cytokines such as IL- 1*β*, TNF*α*, that recruits more leukocytes. Thus, GC may have potential as a dual-purpose preventative and therapeutic with immunostimulatory properties. Future studies are needed to investigate the impact of GC on SARS-CoV-2-specific immune responses.

In addition to the immunoadjuvant properties of GC, direct interactions between GC and SARS-CoV-2 in the nasal cavity may contribute to the protection we observed in mice. As the nasopharyngeal and oral cavity are often the primary entry sites of SARS-CoV-2 and other respiratory viruses, interventions or medications at these locations could provide an important boost in the first line of defense. The reduction in SARS-CoV-2 levels in GC treated mice in the first 2 days post-inoculation suggests that GC has early effects that hamper the initial phase of viral infection, especially given that serum neutralizing antibodies are not detectable before about 7 days. Although we observed that GC reduced tracheal swab titers, we acknowledge that tracheal swabs may less accurately represent virus levels in the upper respiratory tract compared to nasal turbinates. Additionally, GC is viscous, and its cationic charge allows it to bind to negatively charged biomolecules (such as mucin) and act as a mucoadhesive. Our labeling studies with FITC-GC show it remains in the nasal cavity after intranasal administration. GC present at the time of infection may therefore act as a physical barrier and/or directly inactivate SARS-CoV-2 before it can initiate infection. Additional studies of the temporal kinetics of GC bioavailability and detailed studies of GC-virus interactions to establish whether GC blocks virus entry and/or replication are needed to establish mechanisms of protection; these experiments could also clarify why greater protection from lethal disease was observed after inoculation with the higher SARS-CoV-2 dose in these studies.

In summary, we show here that GC treatment of humanized mice pre- and post-infection with SARS-CoV-2 reduces lethal disease, virus levels in the upper respiratory tract, and significant lesions in the lungs. These data suggest a possible role of GC as a SARS-CoV-2 countermeasure. Additional pre-clinical studies could focus on the mechanism of protection, including further assessment of the antiviral and immunomodulatory effects of GC. Future studies should also determine optimal GC dose and schedule that would confer the greatest benefit in reducing disease, as well as to determine whether GC confers both prophylactic and therapeutic benefits in mice that are euthanized at pre-determined intervals instead of when they meet euthanasia criteria.

## METHODS

### Ethics Statement

Mouse work was conducted on protocols #21868 (University of California, Davis), # R20-026 (University of Oklahoma) and # AUP007 (Immunophotonics). Each institutional animal care and use committee (IACUC) approved all mouse work. Infectious virus was handled in certified animal biosafety level 3 laboratory (ABSL-3) spaces in compliance with approved institutional biological use authorization #R2813. The University of California, Davis, is accredited by the Association for Assessment and Accreditation of Laboratory Animal Care (AAALAC). All mouse work adhered to the NIH Guide for the Care and Use of Laboratory Animals[71].

### Mice

Equal numbers of male and female transgenic mice expressing the human ACE2 receptor on a K18 transgene in a C57Bl/6J background (B6.Cg-Tg(K18-ACE2)2Prlmn/J, referenced as ‘hACE2’) were purchased at 5 weeks of age from Jackson Laboratories (Sacramento, CA) and acclimated for up to 6 days. C57BL/6 mice treated with fluorescent GC were used at 6-8 weeks of age and were purchased from Jackson Laboratories. Mice for SARS-CoV-2 studies were co-housed by sex in ABSL-3 conditions. At all 3 institutions, mice were housed in groups of up to 4 animals per cage at 22-25°C and a 12:12 hour light: dark cycle. Rodent chow with 18% protein content and sterile bottled water was provided *ad libitum* for the duration of the experiment.

### Virus

SARS-CoV-2/human/USA/CA-CZB-59X002/2020 (GenBank #MT394528), which was isolated from a patient in 2020 in Northern California and passaged once in Vero- E6 cells, was generously provided by Dr. Christopher Miller (University of California, Davis). To generate stocks for these studies, SARS-CoV-2 was passaged one additional time in Vero-E6 cells to achieve a titer of 2.2 x 107 plaque forming units (PFU)/mL. Single-use virus aliquots were stored at -80°C.

### N-dihydrogalactochitosan Treatment

Sterile 1% weight/volume N-dihydrogalactochitosan (GC) was provided by Immunophotonics. GC was generated using Good Manufacturing Practices (GMP). Testing of GC included appearance, identity (1H NMR and UV/Vis), assay (HPLC), degree of galactation (1H NMR), viscosity, specific gravity, pH, microbiological (endotoxins and sterility), subvisible particulate matter, impurities (boron, galactose, galactitol, transition metals), molecular weight, and polydispersity indices. GC was presented as a 1.0% sterile solution (10 mg/ml) in 5 ml sealed vials and was diluted with sterile deionized water and sterile filtered 20X phosphate buffered saline (PBS) to a final concentration of 0.75% GC and 1X PBS. Mice were anesthetized with isoflurane and 40 µL of either diluted GC or PBS delivery vehicle was administered intranasally (i.n.) by a hanging drop over both nares. Mice were treated identically at 3 days and 1 day prior to inoculation and 1 day post inoculation. Six-week-old male (N=7) and female (N=7) mice were used to assess the histopathologic effects of GC on the murine lung. Mice were GC treated on the same schedule as above, and a subset were euthanized 2 or 14 days after the last GC treatment. Cumulative lung lesion scores were determined using a scoring scale (**Supplementary Table 2**).

### SARS-CoV-2 Inoculation

At inoculation, mice were anesthetized and administered 30 µL of PBS or SARS-CoV-2 diluted in PBS at a dose of 103 or 104 PFU i.n. via hanging drop. Inocula were back-titrated to confirm the target dose. Mice were monitored twice daily for changes in weight, ruffled fur, ataxia, and labored breathing for up to 14 days. On days 1, 2 and 3, mice were anesthetized with isoflurane and throats were swabbed with rayon-tipped swabs (Puritan, Fisher Scientific, Fisher Scientific, Waltham, MA). The process for swabbing was constant across all treatment groups. Swabs were vortexed briefly in 400 µL of Dulbecco’s Modified Eagles Medium (DMEM, Fisher Scientific, Waltham, MA) and frozen at -80°C. Mice were euthanized prior to experimental endpoint if weight loss exceeded 20% of the starting weight or if animals were deemed moribund as evidenced by limb weakness, ataxia or dragging of limbs, loss of limb function or rapid or depressed respiration rate. An adverse event was defined as any moribund disease signs at any time over the duration of the experiment. Prior to euthanasia, whole blood was collected by submandibular vein puncture under isoflurane anesthesia. Whole blood was clotted for >10 min at room temperature then centrifuged for 5 minutes at 8,000 x g and cleared serum was stored at -80°C. Mice were euthanized by isoflurane overdose and cervical dislocation then perfused with cold sterile PBS. Lung (right inferior lobe) and brain (left hemisphere) were weighed and homogenized in 1-10 µL/mg DMEM with a sterile glass bead at 30 Hz for 4 minutes using a TissueLyser (Qiagen, Germantown, MD) automated homogenizer. Homogenates were cleared by centrifugation at 10,000 x g for 4 minutes and the cleared fraction was stored at -80°C.

### Histopathology

At necropsy, lungs were inflated with 10% buffered formalin (Fisher Scientific, Waltham, MA) and mice were fixed for 48 hours at room temperature in a 10- fold volume of 10% buffered formalin. Skulls were demineralized in a 10-fold volume of 0.5 M ethylenediamine tetraacetic acid (EDTA) (pH=7) at 4°C for 14 days, with EDTA solution exchanges every 3 days. Tissues were embedded in paraffin, thin-sectioned, and stained with hematoxylin and eosin (H&E). H&E-stained slides were scanned by a whole-slide image technique using an Aperio slide scanner (Leica, Buffalo Grove, IL) with a resolution of 0.24 μm/pixel. Image files were uploaded on a Leica hosted web- based site and a board certified veterinary anatomic pathologist without knowledge of treatment conditions evaluated sections for SARS-CoV-2 induced histologic lesions. For quantitative assessment of lung inflammation, digital images were captured and analyzed using ImageJ software (Fiji, NIH) to estimate the area of inflamed tissue that was visible to the naked eye at subgross magnification as a percentage of the total surface area of the lung section. Each lung section was scored as described (**Supplementary Table 2**).

### Fluorescent labeled tracing of GC in the nasal cavity

FITC-labeled GC at a concentration of 0.75% (7.5 mg/ml) was applied i.n. in 40 µL to female and male wildtype C57BL/6 mice aged 6-8 weeks (Jackson Laboratories). FITC-GC was manufactured by Immunophotonics, LLC. At 2h, 9h, 2d and 3d (where the last group was administered FITC-GC one additional time 1d before euthanasia) after application, 2 mice per time were euthanized. Two additional mice were not FITC-GC treated; these served as negative controls. The anterior half of the cranium was harvested as described[72] and placed into 4% paraformaldehyde (Cat. 157-8-100, Electron Microscopy Sciences) for 24 h at room temperature. Samples were sent to the St. Louis University (SLU) histology core for processing. The samples were decalcified in ImmunoCal for 2 weeks with the solution changed every 3 days. Next they were washed with PBS then processed into paraffin and embedded. The sections were taken at 4.5 µm every 200 µm. The stained slides were scanned at 20x on an Olympus VS120 to detect the FITC signal. For the unstained slides that were scanned, they first were deparaffinized, then dehydrated so they could be coverslipped and scanned, and 5 µm sections were taken every 200 µm and placed on a slide for examination under an epifluorescence microscope to detect the FITC signal. Nasal concha were evaluated histopathologically and measurements of fluorescence were performed using ImageJ (Fiji). Briefly, images were standardized using the FITC Green channel with rendering set to 1500 for optimal contrast. Screenshots of each micrograph were taken, and the total area of nasal cavity and mucosa was traced and recorded. Teeth, oral mucosa, and connective tissues were excluded for consistency, as sections were variably cut. Fluorescent foci were traced and recorded. Percent area affected was calculated for each slide as total fluorescent area divided by total nasal cavity area. Two sets of measurements were taken per individual, with 4 total observations per treatment since 2 mice were included in each treatment group.

### Plaque Assay

Washes from tracheal swabs, serum, residual inocula, and lung and brain homogenates were thawed and assayed. Samples were serially diluted 10-fold in DMEM with 1% bovine serum albumen (BSA) starting at an initial dilution of 1:8. 125 µL of each dilution was added to confluent Vero CCL-81 cells (ATCC, Manassas, VA) in 12-well plates with cell culture media decanted. Virus was incubated on cells for 1 hour at 5% CO2 in a humidified 37°C incubator. Cell monolayers were overlaid with 0.5% agarose dissolved in DMEM with 5% fetal bovine serum (FBS) and 1x antibiotic-antimycotic (Fisher Scientific, Waltham, MA) and incubated for 3 days at 5% CO2 and 37°C in a humidified incubator. Cells were fixed for >30 minutes with 4% formaldehyde then agarose plugs were removed. Cells were stained with 0.05% crystal violet in 20% ethanol for 10 minutes then rinsed three times with water. Plates were inverted to dry completely and the number of plaques in each well was counted. Viral titers were recorded as the reciprocal of the highest dilution where plaques were noted and are represented as PFU per swab or PFU per mg of solid tissue.

### Plaque Reduction Neutralization Test

Serum collected from mice at day 14 post inoculation was thawed at 37°C and 30 µL was heated in a water bath for 30 minutes at 56°C to inactivate complement proteins. Serum was diluted 4-fold with virus diluent consisting of PBS and 1% FBS, then samples were serially 2-fold diluted 11 times for a dynamic range of 1:4 to 1:4096. An equal volume of virus diluent containing 80 PFU of SARS-CoV-2 was added to each antibody dilution and a no-antibody control consisting of virus diluent only, resulting in a final dynamic range of 1:4 to 1:8192 with one no-antibody control. Antibody-virus dilution series were incubated for 1 hour at 37°C after which they were applied to confluent Vero CCL-81 cells in single-replicate and incubated for 1 hour at 5% CO2 and 37°C in a humidified incubator. Cells were overlaid, incubated, fixed, and stained as described above for plaque assays. Neutralizing titer is defined as the reciprocal of the dilution for which fewer than 20% of plaques were detected versus the no-antibody control (>80% neutralization).

### Statistics

All statistical tests were performed with GraphPad PRISM 9.0.2 (GraphPad Software). Mann Whitney tests were used to compare cumulative pulmonary histologic lesion scores in GC or PBS treated mice. Logrank (Mantel-Cox) test for survival proportions were performed pairwise and *p*-values were adjusted with Bonferroni correction using R version 4.0.0 (R Project) p.adjust function. The correlation between mortality and positive virus detection was calculated by Fisher’s exact test. Repeated measures two-way ANOVA tests were performed on log10-transformed viral titers and multiple comparisons were computed according to Tukey method. Main effect two-way ANOVA tests were performed on mouse weights normalized to starting values at the time of virus inoculation or log10-transformed viral titers and multiple comparisons were computed with Tukey’s method. Area under the curve (AUC) was calculated for longitudinally collected tracheal swabs from days 1,2 and 3 and log10-transformed.

ANOVA of grouped log10-AUC was performed with multiple comparisons computed with Tukey’s method. ANOVA was performed on untransformed histologic scores or percentage of lung affected by inflammation and multiple comparisons were computed with Bonferroni’s method. A Kruskal-Wallis H test was performed on untransformed PRNT80 neutralization values and multiple comparisons were computed according to Dunn’s method.

## DATA AVAILABILITY

All data contributing to the generation of figures and analyses described herein are available at 10.5281/zenodo.7709464.

## AUTHOR CONTRIBUTIONS

Conceptualization: RML, LLC, TH, SSKL, CMW, HL, ARH, and EEB. Investigation: CMW, HL, EEB, ARH, TSW, CFW, EEV. Writing draft: CMW. Review and editing: CMW, HL, EEB, LLC, TH, SSKL, MKK, RML, ARH, TSW, WRC. Visualization: CMW,

EEB, MKK, TSW, LLC. Supervision and project administration: LLC, WRC. Funding acquisition: RML, LLC, WRC.

## ACKNOWLEDGEMENTS

Funding support was provided by the UC Berkeley Henry Wheeler Center for Emerging and Neglected Diseases (CEND) COVID Catalyst Fund, and internal funding from the UC Davis Office of the Vice Chancellor of Research. Funding sources did not influence experimental design and analysis/interpretation of results or impact the decision to publish.

---

**Supplementary Fig. 1.**
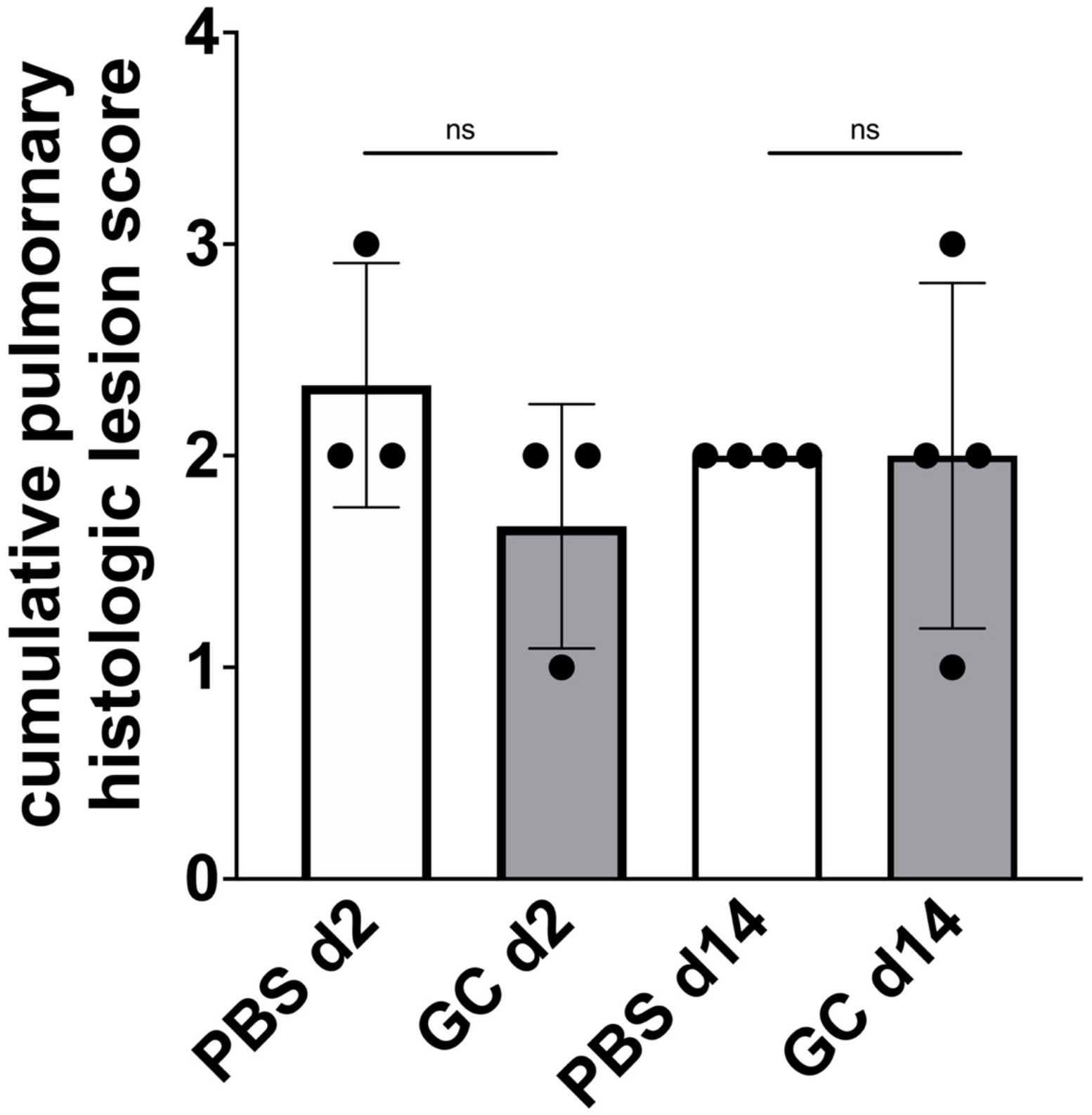
GC administered intranasally does not produce histopathologic changes in murine lung tissues. Six-week-old male and female mice were treated intranasally at 3 intervals with GC or PBS following the same schedule used for mice in the SARS-CoV-2 cohorts. Mice were euthanized 2 or 14 days after the last treatment. Cumulative lung lesion scores were determined using the scoring scale in Supplementary Table 2. A comprehensive list of lesions detected in each animal is shown in Supplementary Table 2. ns is not significant, Mann Whitney test (P=0.6).

**Supplementary Fig. 2:**
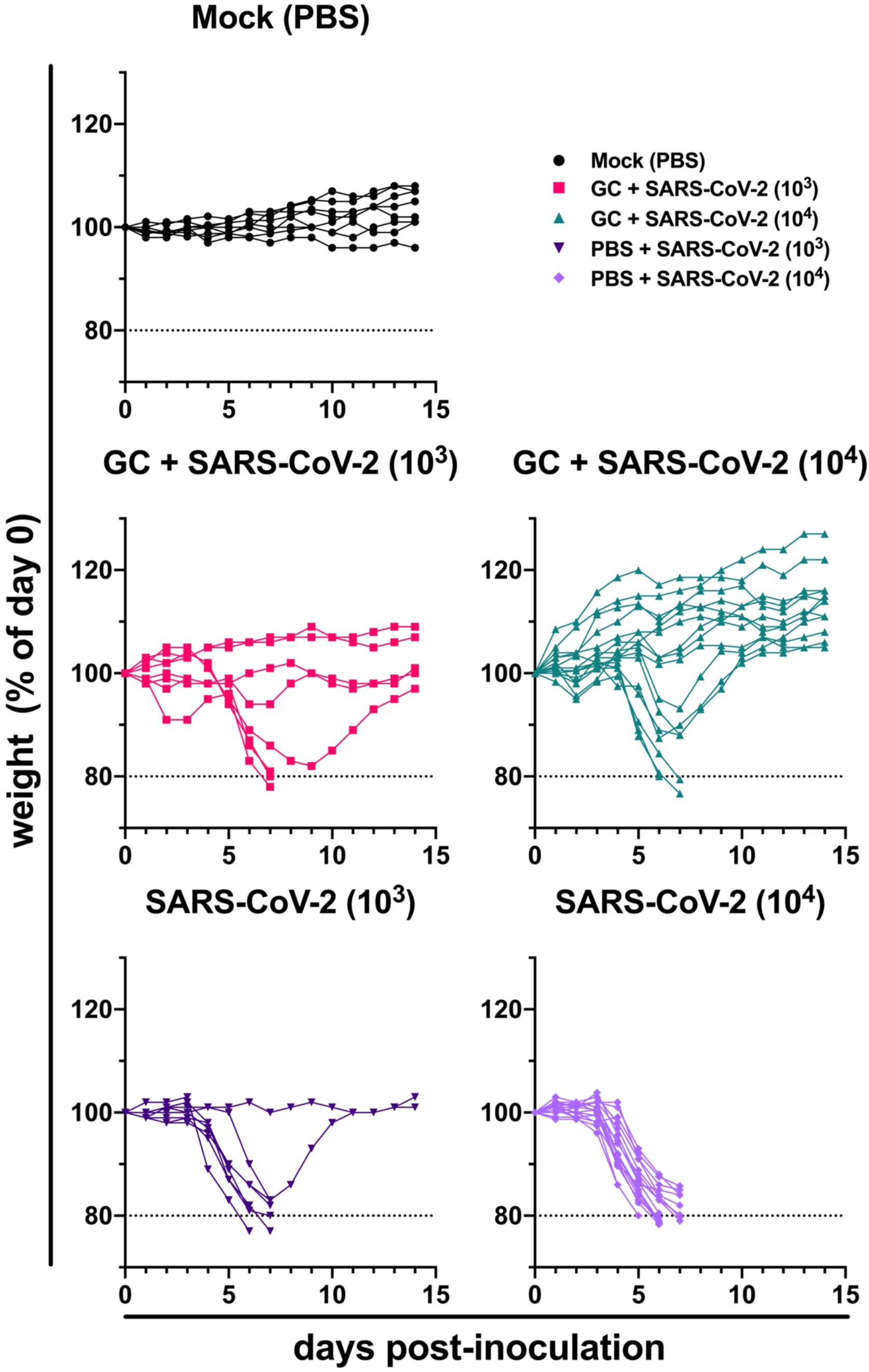
N-dihydrogalactochitosan protects mice from SARS-CoV-2 weight loss. Each line shows individual mouse weight as a percentage of their starting weight at the time of inoculation.

**Supplementary Table 2:** Lung histopathology scoring criteria.

| Score |  | Description |
| --- | --- | --- |
| 0 | none | Within normal limits or rare, scattered lymphocytic infiltrates not observed in control animals but significance questionable (could be background lesion or variation of normal). |
| 1 | minimal | Minimal mononuclear inflammation affecting less than 2% of the section. Inflammatory leukocyte infiltration is limited to a perivascular and/or peribronchiolar distribution with no evidence of alveolar or vascular injury. |
| 2 | mild | Mild peribronchiolar/perivascular inflammation which may also expand alveoli/alveolar septa, composed primarily of macrophages and lymphocytes (+/- scattered neutrophils); increased alveolar macrophages; alveolar septal architecture largely intact; affects 2-10% of the section; and/or scattered alveolar hemorrhage/fibrin/edema; and/or scattered atypical/multinucleated syncytial cells. |
| 3 | moderate | Moderate bronchointerstitial and perivascular inflammation, increased alveolar macrophages, and/or alveolar hemorrhage/fibrin/edema (characterized as above); and/or alveolar damage (characterized by type I pneumocyte necrosis or loss with replacement by hyaline membranes, fibrin, edema, and/or necrotic debris); and/or reparative/regenerative changes (type II pneumocyte hyperplasia, atypical/multinucleated syncytial cells, or fibrosis); lesions affect 10-25% of the section. |
| 4 | severe | As above but more widespread inflammation, hemorrhage/fibrin/ edema, and/or alveolar damage/loss of normal septal architecture; and or regenerative changes affecting greater than 25% of the section. |
| +1 |  | Add 1 point if: Greater than 25% of inflammatory cells are neutrophils; there is significant necrotizing vasculitis, endotheliitis or microthrombi; or if there is |
|  |  | significant bronchiolitis, airway epithelial necrosis or hyperplasia. |

**Supplementary Table 3:** Nasal concha histopathology scoring criteria.

| Score | Degree | Nasal Cavity |
| --- | --- | --- |
| **0** | none | Within normal limits or rare, scattered lymphocytic infiltrates and/or rare foci of mucus, proteinaceous fluid, sloughed epithelium and/or free RBCs not observed in control animals but significance questionable (could be background lesion or variation of normal) |
| **1** | minimal | As above with minimal mononuclear submucosal and/or perivascular inflammation |
| **2** | mild | Mild rhinitis characterized by mucosal/submucosal primarily mononuclear inflammation (+/- low numbers of neutrophils) which may extend into the nasal cavity (along with sloughed epithelial cells and small amounts of mucus/fibrin); overall normal/intact nasal architecture. |
| **3** | moderate | Moderate rhinitis characterized by a mixed inflammatory mucosal/submucosal infiltrate and/or nasal cavity exudate composed of neutrophils, macrophages, and lymphocytes +/- mild to moderate hemorrhage, fibrin, edema and/or necrotic debris. |
| **4** | severe | As above but with more widespread and severe inflammation and/or mucosal erosion/necrosis. |

## REFERENCES

1. Huang C, Wang Y, Li X, Ren L, Zhao J, Hu Y, et al. Clinical features of patients infected with 2019 novel coronavirus in Wuhan, China. The Lancet. 2020. doi:10.1016/S0140-6736(20)30183-5

2. COVID-19 Map - Johns Hopkins Coronavirus Resource Center. [cited 6 Mar 2023]. Available: https://coronavirus.jhu.edu/map.html

3. Wang D, Hu B, Hu C, Zhu F, Liu X, Zhang J, et al. Clinical Characteristics of 138 Hospitalized Patients with 2019 Novel Coronavirus-Infected Pneumonia in Wuhan, China. JAMA - Journal of the American Medical Association. 2020;323: 1061–1069. doi:10.1001/jama.2020.1585

4. Chen N, Zhou M, Dong X, Qu J, Gong F, Han Y, et al. Epidemiological and clinical characteristics of 99 cases of 2019 novel coronavirus pneumonia in Wuhan, China: a descriptive study. The Lancet. 2020;395: 507–513. doi:10.1016/S0140-6736(20)30211-7

5. Zhou F, Yu T, Du R, Fan G, Liu Y, Liu Z, et al. Clinical course and risk factors for mortality of adult inpatients with COVID-19 in Wuhan, China: a retrospective cohort study. The Lancet. 2020;395: 1054–1062. doi:10.1016/S0140-6736(20)30566-3

6. Hellmuth J, Barnett TA, Asken BM, Kelly JD, Torres L, Stephens ML, et al. Persistent COVID-19-associated neurocognitive symptoms in non-hospitalized patients. J Neurovirol. 2021;27: 191–195. doi:10.1007/s13365-021-00954-4

7. Graham EL, Clark JR, Orban ZS, Lim PH, Szymanski AL, Taylor C, et al. Persistent neurologic symptoms and cognitive dysfunction in non-hospitalized Covid-19 “long haulers.” Ann Clin Transl Neurol. 2021. doi:10.1002/acn3.51350

8. Lu R, Zhao X, Li J, Niu P, Yang B, Wu H, et al. Genomic characterisation and epidemiology of 2019 novel coronavirus: implications for virus origins and receptor binding. The Lancet. 2020;395: 565–574. doi:10.1016/S0140-6736(20)30251-8

9. Zhou P, Yang X Lou, Wang XG, Hu B, Zhang L, Zhang W, et al. A pneumonia outbreak associated with a new coronavirus of probable bat origin. Nature. 2020;579: 270–273. doi:10.1038/s41586-020-2012-7

10. Lakdawala SS, Menachery VD. The search for a COVID-19 animal model. Science (1979). 2020;368: 942–943. doi:10.1126/science.abc6141

11. Muñoz-Fontela C, Dowling WE, Funnell SGP, Gsell PS, Riveros-Balta AX, Albrecht RA, et al. Animal models for COVID-19. Nature. 2020;586: 509– 515. doi:10.1038/s41586-020-2787-6

12. McCray PB, Pewe L, Wohlford-Lenane C, Hickey M, Manzel L, Shi L, et al. Lethal Infection of K18-hACE2 Mice Infected with Severe Acute Respiratory Syndrome Coronavirus. J Virol. 2007. doi:10.1128/jvi.02012-06

13. Winkler ES, Bailey AL, Kafai NM, Nair S, McCune BT, Yu J, et al. SARS-CoV-2 infection of human ACE2-transgenic mice causes severe lung inflammation and impaired function. Nat Immunol. 2020;21: 1327–1335. doi:10.1038/s41590-020-0778-2

14. Zheng J, Wong LYR, Li K, Verma AK, Ortiz ME, Wohlford-Lenane C, et al. COVID-19 treatments and pathogenesis including anosmia in K18-hACE2 mice. Nature. 2021;589: 603–607. doi:10.1038/s41586-020-2943-z

15. Symptoms of COVID-19 | CDC. [cited 21 May 2021]. Available: https://www.cdc.gov/coronavirus/2019-ncov/symptoms-testing/symptoms.html

16. FDA Issues Emergency Use Authorization for Third COVID-19 Vaccine | FDA. [cited 26 Apr 2021]. Available: https://www.fda.gov/news-events/press-announcements/fda-issues-emergency-use-authorization-third-covid-19-vaccine

17. Coronavirus disease (COVID-19): Herd immunity, lockdowns and COVID- 19. [cited 26 Apr 2021]. Available: https://www.who.int/news-room/q-a-detail/herd-immunity-lockdowns-and-covid-19

18. Rhoades A. Veklury (remdesivir) EUA Letter of Approval, reissued 10/22/2020. 2020.

19. Beigel JH, Tomashek KM, Dodd LE, Mehta AK, Zingman BS, Kalil AC, et al. Remdesivir for the Treatment of Covid-19 — Final Report. New England Journal of Medicine. 2020;383: 1813–1826. doi:10.1056/NEJMoa2007764

20. WHO recommends against the use of remdesivir in COVID-19 patients. [cited 25 Apr 2021]. Available: https://www.who.int/news-room/feature-stories/detail/who-recommends-against-the-use-of-remdesivir-in-covid-19-patients

21. Administration D. FDA Combating COVID-19 With Therapeutics. 2020.

22. Emergency Use Authorization | FDA. [cited 7 Mar 2023]. Available: https://www.fda.gov/emergency-preparedness-and-response/mcm-legal-regulatory-and-policy-framework/emergency-use-authorization#coviddrugs

23. Chirkov SN. The antiviral activity of chitosan (review). Applied Biochemistry and Microbiology. 2002. pp. 1–8. doi:10.1023/A:1013206517442

24. Mehrabi M, Montazeri H, Dounighi MN, Rashti A, Vakili-Ghartavol R. Chitosan-based nanoparticles in mucosal vaccine delivery. Archives of Razi Institute. Razi Vaccine and Serum Research Institute; 2018. pp. 165–176. doi:10.22092/ari.2017.109235.1101

25. Li X, Min M, Du N, Gu Y, Hode T, Naylor M, et al. Chitin, chitosan, and glycated chitosan regulate immune responses: The novel adjuvants for cancer vaccine. Clinical and Developmental Immunology. Hindawi Limited; 2013. doi:10.1155/2013/387023

26. Mudgal J, Mudgal PP, Kinra M, Raval R. Immunomodulatory role of chitosan-based nanoparticles and oligosaccharides in cyclophosphamide-treated mice. Scand J Immunol. 2019;89: e12749. doi:10.1111/sji.12749

27. Carroll TD, Jegaskanda S, Matzinger SR, Fritts L, McChesney MB, Kent SJ, et al. A lipid/DNA adjuvant-inactivated influenza virus vaccine protects rhesus macaques from uncontrolled virus replication afer heterosubtypic influenza a virus challenge. Journal of Infectious Diseases. 2018. doi:10.1093/infdis/jiy238

28. Lei X, Dong X, Ma R, Wang W, Xiao X, Tian Z, et al. Activation and evasion of type I interferon responses by SARS-CoV-2. Nat Commun. 2020;11. doi:10.1038/s41467-020-17665-9

29. Xia H, Cao Z, Xie X, Zhang X, Chen JYC, Wang H, et al. Evasion of Type I Interferon by SARS-CoV-2. Cell Rep. 2020;33. doi:10.1016/j.celrep.2020.108234

30. Ribero MS, Jouvenet N, Dreux M, Nisole S. Interplay between SARS-CoV-2 and the type I interferon response. PLoS Pathogens. Public Library of Science; 2020. doi:10.1371/journal.ppat.1008737

31. Zheng M, Qu D, Wang H, Sun Z, Liu X, Chen J, et al. Intranasal Administration of Chitosan Against Influenza A (H7N9) Virus Infection in a Mouse Model. Sci Rep. 2016;6. doi:10.1038/srep28729

32. Wang X, Zhang W, Liu F, Zheng M, Zheng D, Zhang T, et al. Intranasal immunization with live attenuated influenza vaccine plus chitosan as an adjuvant protects mice against homologous and heterologous virus challenge. Arch Virol. 2012;157: 1451–1461. doi:10.1007/s00705-012-1318-7

33. Choi B, Jo DH, Anower AKMM, Islam SMS, Sohn S. Chitosan as an Immunomodulating Adjuvant on T-Cells and Antigen-Presenting Cells in Herpes Simplex Virus Type 1 Infection. Mediators Inflamm. 2016;2016. doi:10.1155/2016/4374375

34. Pyrć K, Milewska A, Duran EB, Botwina P, Dabrowska A, Jedrysik M, et al. SARS-CoV-2 inhibition using a mucoadhesive, amphiphilic chitosan that may serve as an anti-viral nasal spray. Sci Rep. 2021;11. doi:10.1038/s41598-021-99404-8

35. Jaber N, Al-Remawi M, Al-Akayleh F, Al-Muhtaseb N, Al-Adham ISI, Collier PJ. A review of the antiviral activity of Chitosan, including patented applications and its potential use against COVID-19. J Appl Microbiol. 2022;132: 41–58. doi:10.1111/jam.15202

36. Safarzadeh M, Sadeghi S, Azizi M, Rastegari-Pouyani M, Pouriran R, Haji Molla Hoseini M. Chitin and chitosan as tools to combat COVID-19: A triple approach. Int J Biol Macromol. 2021;183: 235–244. doi:10.1016/j.ijbiomac.2021.04.157

37. Sharma N, Modak C, Singh PK, Kumar R, Khatri D, Singh SB. Underscoring the immense potential of chitosan in fighting a wide spectrum of viruses: A plausible molecule against SARS-CoV-2? Int J Biol Macromol. 2021;179: 33–44. doi:10.1016/j.ijbiomac.2021.02.090

38. Zhuo S-H, Wu J-J, Zhao L, Li W-H, Zhao Y-F, Li Y-M. A chitosan-mediated inhalable nanovaccine against SARS-CoV-2. Nano Res. 2022;15: 4191– 4200. doi:10.1007/s12274-021-4012-9

39. Tan RSL, Hassandarvish P, Chee CF, Chan LW, Wong TW. Chitosan and its derivatives as polymeric anti-viral therapeutics and potential anti-SARS-CoV-2 nanomedicine. Carbohydr Polym. 2022;290: 119500. doi:10.1016/j.carbpol.2022.119500

40. Chen WR, Carubelli R, Liu H, Nordquist RE. Laser immunotherapy: A novel treatment modality for metastatic tumors. Applied Biochemistry and Biotechnology - Part B Molecular Biotechnology. 2003;25: 37–43. doi:10.1385/MB:25:1:37

41. Song S, Zhou F, Nordquist RE, Carubelli R, Liu H, Chen WR. Glycated chitosan as a new non-toxic immunological stimulant Glycated chitosan immunological stimulant. Immunopharmacol Immunotoxicol. 2009;31: 202– 208. doi:10.1080/08923970802629593

42. Zizzari I, Napoletano C, Battisti F, Rahimi H, Caponnetto S, Pierelli L, et al. MGL Receptor and Immunity: When the Ligand Can Make the Difference. J Immunol Res. 2015;2015. doi:10.1155/2015/450695

43. Korbelik M, Banáth J, Zhang W, Hode T, Lam S, Gallagher P, et al. N-dihydrogalactochitosan-supported tumor control by photothermal therapy and photothermal therapy-generated vaccine. J Photochem Photobiol B. 2020;204. doi:10.1016/J.JPHOTOBIOL.2020.111780

44. Korbelik M, Hode T, Lam SSK, Chen WR. Novel Immune Stimulant Amplifies Direct Tumoricidal Effect of Cancer Ablation Therapies and Their Systemic Antitumor Immune Efficacy. Cells. NLM (Medline); 2021. doi:10.3390/cells10030492

45. Qi S, Lu L, Zhou F, Chen Y, Xu M, Chen L, et al. Neutrophil infiltration and whole-cell vaccine elicited by N-dihydrogalactochitosan combined with NIR phototherapy to enhance antitumor immune response and T cell immune memory. Theranostics. 2020;10: 1814–1832. doi:10.7150/thno.38515

46. Zhou F, Yang J, Zhang Y, Liu M, Lang ML, Li M, et al. Local phototherapy synergizes with immunoadjuvant for treatment of pancreatic cancer through induced immunogenic tumor vaccine. Clinical Cancer Research. 2018;24: 5335–5346. doi:10.1158/1078-0432.CCR-18-1126

47. El-Hussein A, Lam SSK, Raker J, Chen WR, Hamblin MR. N-dihydrogalactochitosan as a potent immune activator for dendritic cells. J Biomed Mater Res A. 2017;105: 963–972. doi:10.1002/jbm.a.35991

48. Hoover AR, Liu K, Devette CI, Krawic JR, West CL, Medcalf D, et al. ScRNA-seq reveals tumor microenvironment remodeling induced by local intervention-based immunotherapy. bioRxiv. 2020; 2020.10.02.323006. doi:10.1101/2020.10.02.323006

49. Hoover AR, More S, Liu K, West CL, Valero TI, Yu N, et al. A novel biopolymer for mucosal adjuvant against respiratory pathogens. bioRxiv. 2022; 2022.09.07.506979. doi:10.1101/2022.09.07.506979

50. Brown C, Vostok J, Johnson H, Burns M, Gharpure R, Sami S, et al. Outbreak of SARS-CoV-2 Infections, Including COVID-19 Vaccine Breakthrough Infections, Associated with Large Public Gatherings - Barnstable County, Massachusetts, July 2021. MMWR Morb Mortal Wkly Rep. 2021;70: 1059–1062. doi:10.15585/MMWR.MM7031E2

51. Bellich B, D’Agostino I, Semeraro S, Gamini A, Cesàro A. “The Good, the Bad and the Ugly” of Chitosans. Mar Drugs. 2016;14. doi:10.3390/MD14050099

52. Jayathilakan K, Sultana K, Radhakrishna K, Bawa AS. Utilization of byproducts and waste materials from meat, poultry and fish processing industries: A review. Journal of Food Science and Technology. J Food Sci Technol; 2012. pp. 278–293. doi:10.1007/s13197-011-0290-7

53. Negm NA, Hefni HHH, Abd-Elaal AAA, Badr EA, Abou Kana MTH. Advancement on modification of chitosan biopolymer and its potential applications. International Journal of Biological Macromolecules. Elsevier B.V.; 2020. pp. 681–702. doi:10.1016/j.ijbiomac.2020.02.196

54. Mokhtar H, Biffar L, Somavarapu S, Frossard JP, McGowan S, Pedrera M, et al. Evaluation of hydrophobic chitosan-based particulate formulations of porcine reproductive and respiratory syndrome virus vaccine candidate T cell antigens. Vet Microbiol. 2017;209: 66–74. doi:10.1016/j.vetmic.2017.01.037

55. Shim S, Park HE, Soh SH, Im Y Bin, Yoo HS. Induction of Th2 response through TLR2-mediated MyD88-dependent pathway in human microfold cells stimulated with chitosan nanoparticles loaded with Brucella abortus Mdh. Microb Pathog. 2020;142. doi:10.1016/j.micpath.2020.104040

56. Zhang X, Wu K, Wang D, Yue X, Song D, Zhu Y, et al. Nucleocapsid protein of SARS-CoV activates interleukin-6 expression through cellular transcription factor NF-κB. Virology. 2007;365: 324–335. doi:10.1016/j.virol.2007.04.009

57. Blanco-Melo D, Nilsson-Payant BE, Liu WC, Uhl S, Hoagland D, Møller R, et al. Imbalanced Host Response to SARS-CoV-2 Drives Development of COVID-19. Cell. 2020;181: 1036–1045.e9. doi:10.1016/j.cell.2020.04.026

58. Pyle C, Uwadiae F, Swieboda D, Harker J. Early IL-6 signalling promotes IL-27 dependent maturation of regulatory T cells in the lungs and resolution of viral immunopathology. PLoS Pathog. 2017;13. doi:10.1371/JOURNAL.PPAT.1006640

59. Bueter CL, Lee CK, Rathinam VAK, Healy GJ, Taron CH, Specht CA, et al. Chitosan but not chitin activates the inflammasome by a mechanism dependent upon phagocytosis. Journal of Biological Chemistry. 2011;286: 35447–35455. doi:10.1074/jbc.M111.274936

60. Carroll EC, Jin L, Mori A, Muñoz-Wolf N, Oleszycka E, Moran HBT, et al. The Vaccine Adjuvant Chitosan Promotes Cellular Immunity via DNA Sensor cGAS-STING-Dependent Induction of Type I Interferons. Immunity. 2016;44: 597–608. doi:10.1016/j.immuni.2016.02.004

61. Sun B, Yu S, Zhao D, Guo S, Wang X, Zhao K. Polysaccharides as vaccine adjuvants. Vaccine. Elsevier Ltd; 2018. pp. 5226–5234. doi:10.1016/j.vaccine.2018.07.040

62. Read RC, Naylor SC, Potter CW, Bond J, Jabbal-Gill I, Fisher A, et al. Effective nasal influenza vaccine delivery using chitosan. Vaccine. 2005;23: 4367–4374. doi:10.1016/j.vaccine.2005.04.021

63. Elieh Ali Komi D, Sharma L, Dela Cruz CS. Chitin and Its Effects on Inflammatory and Immune Responses. Clinical Reviews in Allergy and Immunology. Humana Press Inc.; 2018. pp. 213–223. doi:10.1007/s12016-017-8600-0

64. Bashiri S, Koirala P, Toth I, Skwarczynski M. Carbohydrate immune adjuvants in subunit vaccines. Pharmaceutics. MDPI AG; 2020. pp. 1–33. doi:10.3390/pharmaceutics12100965

65. Bueter CL, Specht CA, Levitz SM. Innate Sensing of Chitin and Chitosan. PLoS Pathog. 2013;9. doi:10.1371/journal.ppat.1003080

66. Seetharaman J, Kfanigsberg A, Slaaby R, Leffler H, Barondes SH, Rini JM. X-ray crystal structure of the human galectin-3 carbohydrate recognition domain at 2.1-Å resolution. Journal of Biological Chemistry. 1998;273: 13047–13052. doi:10.1074/jbc.273.21.13047

67. Semeňuk T, Krist P, Pavlíček J, Bezouška K, Kuzma M, Novák P, et al. Synthesis of chitooligomer-based glycoconjugates and their binding to the rat natural killer cell activation receptor NKR-P1. Glycoconj J. 2001;18: 817–826. doi:10.1023/A:1021111703443

68. Zhou F, Wu S, Song S, Chen WR, Resasco DE, Xing D. Antitumor immunologically modified carbon nanotubes for photothermal therapy. Biomaterials. 2012;33: 3235–3242. doi:10.1016/j.biomaterials.2011.12.029

69. Nevagi RJ, Khalil ZG, Hussein WM, Powell J, Batzloff MR, Capon RJ, et al. Polyglutamic acid-trimethyl chitosan-based intranasal peptide nano-vaccine induces potent immune responses against group A streptococcus. Acta Biomater. 2018;80: 278–287. doi:10.1016/j.actbio.2018.09.037

70. Micoli F, Costantino P, Adamo R. Potential targets for next generation antimicrobial glycoconjugate vaccines. FEMS Microbiology Reviews. Oxford University Press; 2018. pp. 388–423. doi:10.1093/femsre/fuy011

71. National Institutes of Health. GUIDE LABORATORY ANIMALS FOR THE CARE AND USE OF Eighth Edition Committee for the Update of the Guide for the Care and Use of Laboratory Animals Institute for Laboratory Animal Research Division on Earth and Life Studies. 2011 [cited 6 Mar 2023]. Available: http://www.nap.edu.

72. Cisney ED, Fernandez S, Hall SI, Krietz GA, Ulrich RG. Examining the role of nasopharyngeal-associated lymphoreticular tissue (NALT) in mouse responses to vaccines. J Vis Exp. 2012 [cited 6 Mar 2023]. doi:10.3791/3960

